# Modulation of high frequency by low-frequency Oscillations in the basal ganglia

**DOI:** 10.1101/196592

**Authors:** D. Nouri, R. Ebrahimpour, A. Mirzaei

**Affiliations:** Department of Computer Engineering, Shahid Beheshti University, Tehran, Iran; School of Cognitive Sciences, Institute for Research in Fundamental Sciences (IPM), Tehran, Iran; Cognitive Science Research Lab., Department of Computer Engineering, Shahid Rajaee Teacher Training University, Tehran, Iran; BrainLinks-BrainTools, University of Freiburg, Freiburg, Germany; Faculty of Biology, University of Freiburg, Freiburg, Germany

**Keywords:** Basal ganglia, cross-frequency coupling, nested oscillations, cortico-basal ganglia network, Parkinson’s disease

## Abstract

Modulation of beta band fioscillatory activity (15-30 Hz) by delta band oscillatory activity (1-3 Hz) in the cortico-basal ganglia loop is important for normal basal ganglia functions. However, the neural mechanisms underlying this modulation are poorly understood. To understand the mechanisms underlying such frequency modulations in the basal ganglia, we use large scale subthalamo-pallidal network model stimulated via a delta-frequency input signal. We show that inhibition of external Globus Pallidus (GPe) and excitation of the Subthalamic nucleus (STN) using the delta-band stimulation leads to the same delta-beta interactions in the network model as the experimental results observed in healthy basal ganglia. In addition, we show that pathological beta oscillations in the network model decorrelates the delta-beta link in the network model. In general, using our simulation results, we propose that striato-pallidal inhibition and cortico-subthalamic excitation are the potential sources of the delta-beta link observed in the intact basal ganglia.

## 1 Introduction

Modulated high-frequency oscillations by low-frequency oscillations in the brain continue to provide important clues about healthy and pathological neuronal activities [1]. More specifically, modulation of beta frequency oscillations (i.e. 15-30 Hz) by delta frequency oscillations (1-3 Hz) is an indication of healthy basal ganglia [2, 3, and 4]. It has been shown that beta power of the Local Field Potential (LFP) data recorded from GPe in the basal ganglia is positively correlated with delta power in the healthy state. However, such positive correlation does not exist in Parkinson’s disease (PD) state [5]. This is accompanied by reduction in information processing capability of the basal ganglia [6, 7, 8, 9, and 10].

The underlying mechanism of such positive correlation between delta-beta band neuronal activities in the healthy basal ganglia and how it is affected in the pathological state are still unknown. Here we show how delta band oscillations of the STN and GPe can modulate beta oscillations in a subthalamo-pallidal network model. Using our simulation results we also show that pathological PD-like beta oscillations in the network model lead to delta-beta decorrelation in the network model, similar to the experimental LFP data [5]

## 2 Materials and methods

### Network model

The network model we use comprises 2000 inhibitory (i.e. GPe) and 1000 excitatory (i.e. STN) neurons. Neurons were implemented as Leaky-Integrate-and-Fire (LIF) neuron type. STN neurons excite GPe neurons while receiving inhibitory input from GPe. STN neurons receive excitatory synaptic inputs from other STN neurons. GPe neurons receive inhibitory synaptic inputs from other GPe neurons. The network model parameters (i.e. synaptic strengths and synaptic probabilities) are given in [1].

### External input

Excitatory uncorrelated spike trains are used as external input to provide baseline activity of each population in the network model. These inputs were used to obtain realistic baseline firing rates in STN (∼15 Hz) and GPe (∼45 Hz), observed in the healthy state [12, 13].

### Striatal stimulation

Uncorrelated Poisson spike trains (representing indirect pathway striatal activity) were used to inhibit GPe neurons. Firing rate of the Poisson spike trains was systematically varied from 0 Hz (healthy state) to 500 Hz (PD state) to generate Fig 5.

### Delta band sinusoidal stimulation

We used inhomogeneous Poisson spike trains with firing rate pattern of a sinusoidal signal of 3 Hz as delta band stimulation of the network model. The delta band stimulation was used to stimulate the network in four different stimulation scenarios: 1) excitation of STN, 2) inhibition of STN, 3) excitation of GPe, and 4) inhibition of GPe.

### Time resolved population firing rate

Population activity (for both STN and GPe) in the network model was computed by counting total number of spikes within a sliding 5 ms time window and dividing the resulting value by (0.005 * number of neurons) to obtain the population firing rate in Hz. The same method was applied for each shifted 5 ms time window (with 1 ms steps) to get the population firing rate over time. To avoid initial network transients, the first 1000 milliseconds was excluded from further analysis.

### Time-frequency analysis

Spectrograms were computed by convolving the STN (or GPe) population firing rate in the network model with a standard Morlet wavelet of integer frequencies (f = 1 to 100 Hz) and taking the logarithm of the squared magnitude of the resulting time series. In order to generate Fig 4A, bottom, the mean spectrogram across 100 simulations of the model was computed.

### Simulation tools

All network simulations and data analysis were written in Python using PyNN as an interface to the simulation environment NEST [14].

## 3 Results

To understand the mechanisms underlying modulation of beta oscillations by delta oscillation in the basal ganglia, we exposed the network model to the delta band frequency stimulation (see Methods for more details). To study how stimulation of the network model in delta frequency band affects the network model dynamics, four different delta stimulation scenarios were considered: 1) excitation of STN, 2) inhibition of STN, 3) excitation of GPe, and 4) inhibition of GPe. Each stimulation scenario was individually investigated in the network model to see which one can lead to the same delta-beta dynamics observed in the experimental data [5].

**Fig 1.**
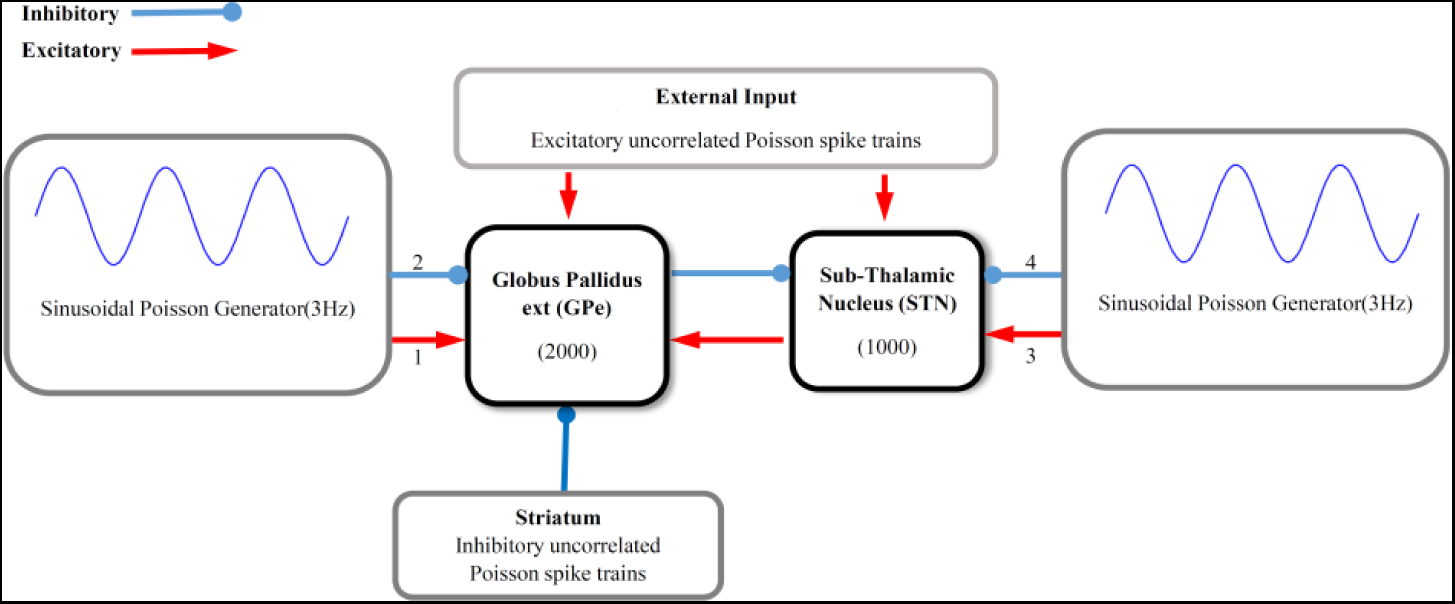
Schematic of the network model exposed to the delta band sinusoidal stimulation pattern. The network model comprises STN and GPe neuronal populations (see Methods for more details). Excitatory uncorrelated spike trains were used as external input to provide baseline activity of each neural population. Uncorrelated Poisson spike trains were used as striatal input to GPe. The strength of GPe inhibition was systematically varied between 0 Hz (healthy state) to 500 Hz (PD state). The network was stimulated via delta band stimulation pattern (see Methods). Four different stimulation scenarios were considered: (1) Excitation of STN (2) Inhibition of STN (3) Excitation of GPe and (4) Inhibition of GPe. All excitatory and inhibitory connections are shown in red and blue, respectively.

### 3.1 Interactions between delta and beta band oscillations in the network model

Our simulation results show that excitation of the STN and inhibition of the GPe in delta frequency band can lead to the beta oscillations in the network model. The network beta oscillations emerged at the certain phase of the delta stimulation pattern (e.g. mainly around the peak; Fig 2A, B). However, when the delta band stimulation was used to excite the GPe or to inhibit the STN, it could not generate beta oscillations in the network model (Fig 2C, D). Therefore, we conclude that the potential mechanism underlying experimentally observed the relationship between delta and beta band LFP oscillations can be delta band excitatory input to STN (i.e. through cortical drive) or inhibitory input to GPe (i.e. through striatal drive).

**Fig 2.**
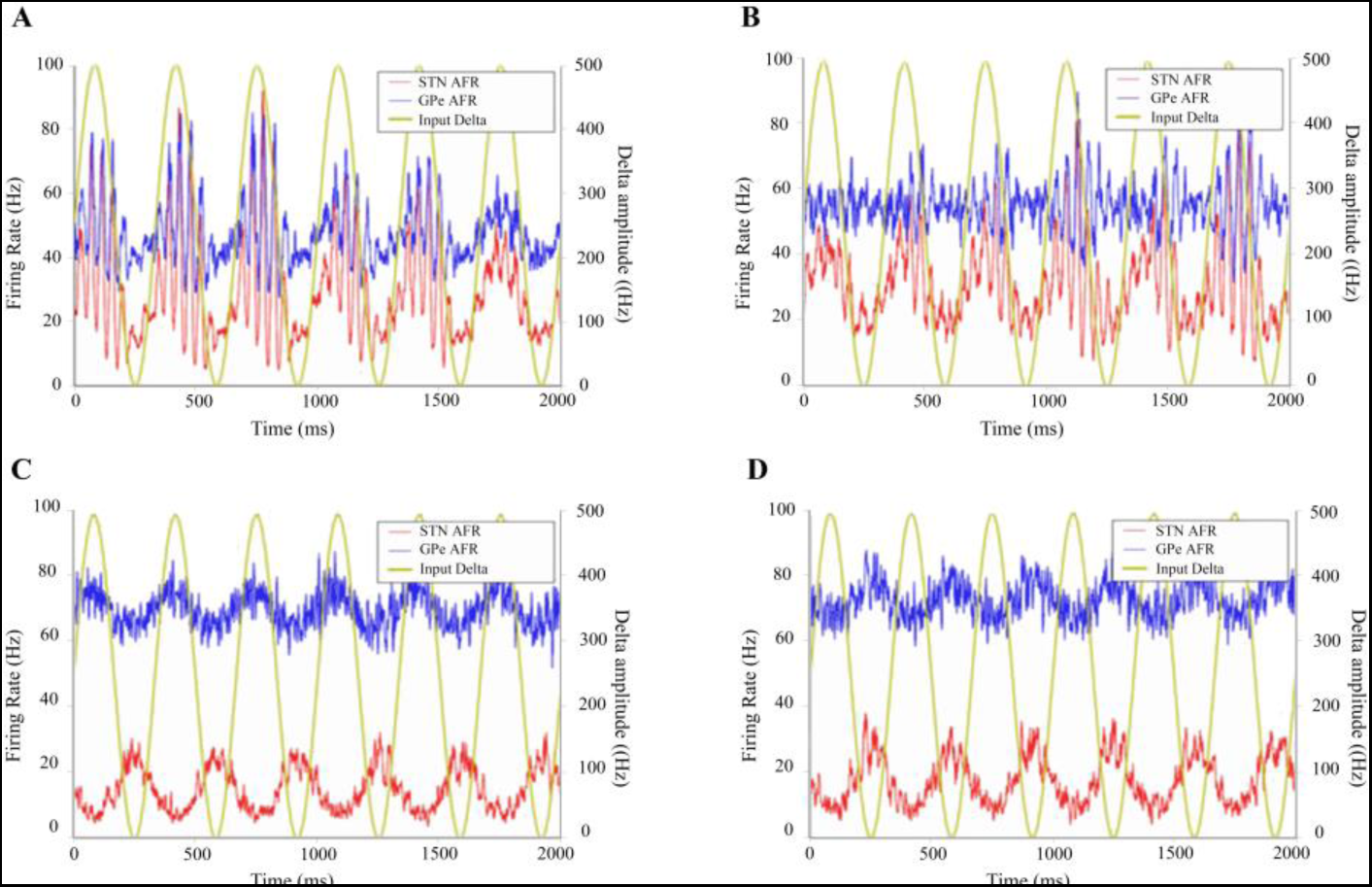
Response of the network model to the delta band sinusoidal stimulation. (A-D) STN and GPe population firing rates (blue and red, respectively) in response to the delta band sinusoidal excitatory input to STN (A), inhibitory input to GPe (B), excitatory input to GPe (C), and inhibitory input to STN (D). The delta band stimulation is shown in yellow.

Computing the spectrograms of the GPe and STN population firing rates reveals the interplay between beta-band and delta-band oscillations. It is obvious from this analysis that maximum level of the beta power occurs at the delta peak. (Fig 4A, B, bottom). Such a relationship between delta and beta band oscillation has been reported in experimental studies in the healthy state which is shown in Fig 3 [5, 15].

**Fig 3.**
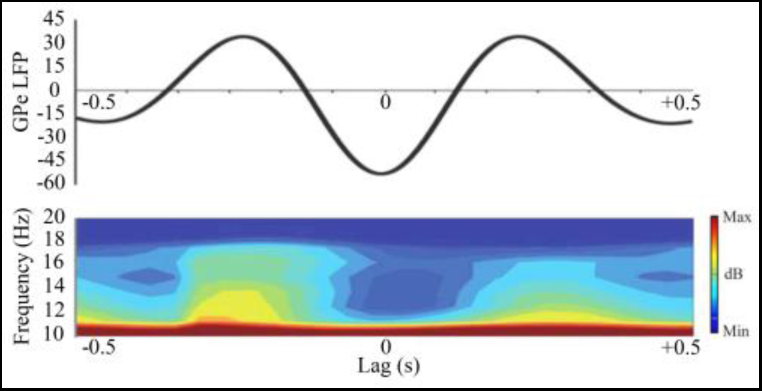
An example of spectra analysis from GPe in the basal ganglia that demonstrates delta-beta relationship (adapted from [5]). The top panel shows the mean delta filtered LFP from one rat. The bottom panel shows modulation of beta power, corresponding to the top panel.

**Fig 4.**
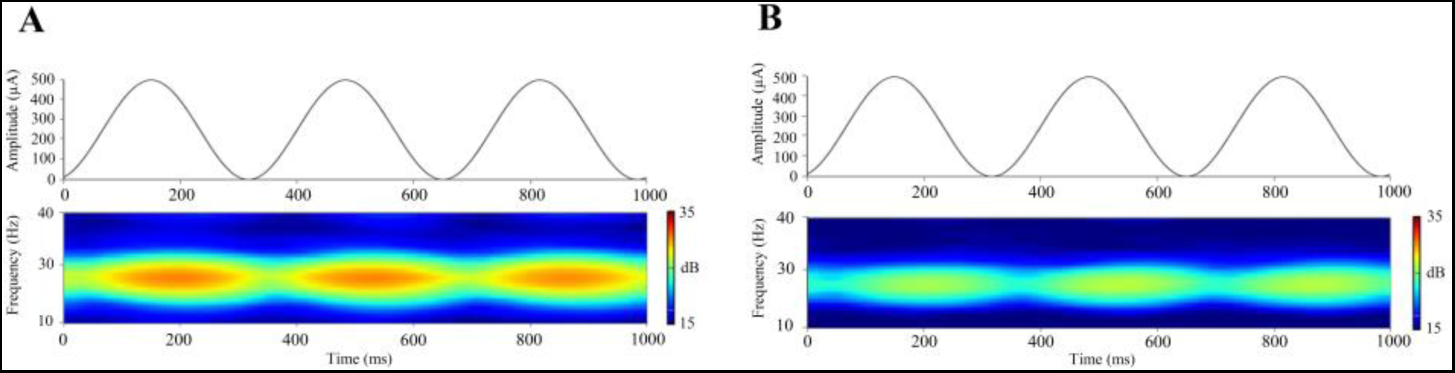
Fluctuations in the beta power due to delta band stimulation in the STN-GPe network. (A) Top, Delta band firing pattern of the inhomogeneous Poisson spike trains are used as excitatory input to STN. Bottom, the spectrogram of the STN population activity in response to excitation of STN using the delta band stimulation (average of 100 simulations). B is same as A, for inhibition of GPe.

### 3.2 Delta-beta relationship disappears in pathological state of the network model

The activity of the indirect striatal units pathologically increases in PD which leads to a strong striato-pallidal inhibition in the basal ganglia [16]. To simulate progress of PD in the network model, we increased the strength of the striato-pallidal inhibition from 0 Hz (healthy state) to 500 Hz (pathological state). It has previously been shown that such strong striato-pallidal inhibition leads to beta oscillations in the network model, similar to what has been observed in the basal ganglia of patients with PD [11]. Using our simulation results, we show that as beta power increases (i.e. during the progress of PD) the delta power decreases, consequently. In other words, strengthening the striatal inhibitory input to GPe in the network model leads to increase in pathological beta power and subsequently decrease in delta power (Fig 5). Therefore, in PD state of the network model the delta-beta relationship does not exist anymore, similar to the experimental results [5].

**Fig 5.**
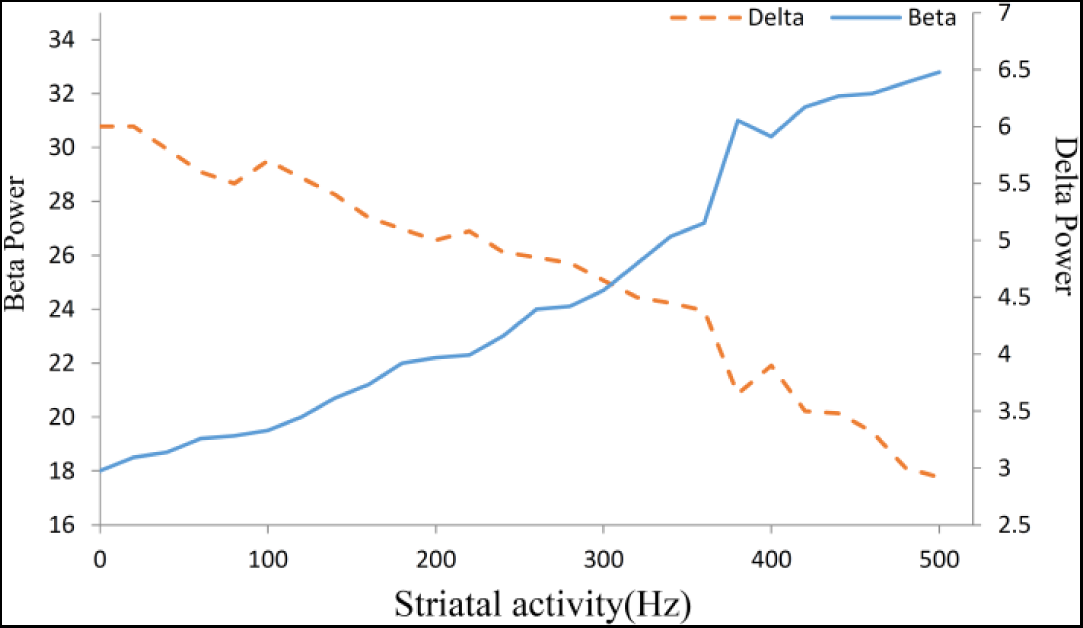
Delta power decreases when the pathological beta power increases. Mean beta power (blue) and mean delta power (red; average of 100 simulations) during the progression of PD in the network model. Note that the striatal activity (x-axis) increases during progression of PD.

## 4 Conclusion

Using our simulation results, we conclude that delta-band excitation of STN (presumably through cortical drive) and inhibition of GPe (presumably through striatal drive) leads to the same relationship between delta and beta band oscillations in the network model as the experimental LFP results. Moreover, similar to the experimental results, we observed in our simulation results that the delta-beta relationship disappears in PD state of the network model.

